# Nitric oxide regulates growth coordination during regeneration

**DOI:** 10.1101/016030

**Authors:** Jacob S. Jaszczak, Jacob B. Wolpe, Anh Q. Dao, Adrian Halme

## Abstract

Mechanisms that coordinate the growth of different tissues during development are essential for producing adult animals with proper organ proportion. Here we describe a pathway through which tissues communicate with each other to coordinate growth. During *Drosophila melanogaster* larval development, damage to imaginal discs activates a regeneration checkpoint that produces both a delay in developmental timing and slows the growth of undamaged tissues, coordinating regeneration of the damaged tissue with developmental progression and overall growth. Both developmental delay and growth control are mediated by secretion of the insulin/relaxin family peptide Dilp8 from regenerating tissues. Here we demonstrate that Dilp8-dependent growth coordination between regenerating and undamaged tissues, but not developmental delay, requires the activity of nitric oxide synthase (NOS) in the prothoracic gland. NOS limits the growth of undamaged tissues by reducing ecdysone biosynthesis, a requirement for imaginal disc growth during both the regenerative checkpoint and normal development. Therefore, NOS activity in the prothoracic gland translates information about the growth status of individual tissues into coordinated tissue growth through the regulation of endocrine signals.

## INTRODUCTION

Allometry, broadly defined as the scaling of organ growth, can have a profound impact on the biological function of animals. There are many examples throughout nature of how allometric growth regulation can influence biological function. For instance, in the horned dung beetle *Onthophagus netriventris* an inverse allometry is observed between horn and testes size in male beetles, producing distinct reproductive strategies (1, 2). Allometric growth regulation can also impact human health, where variation from optimal relative heart size can increase susceptibility for cardiovascular disease (3). Despite the fundamental role of growth scaling in biology, no pathways have been described that explain how the coordination of growth between different tissues is regulated. Experimental understanding of growth regulation has been primarily focused on either tissue-autonomous mechanisms of growth regulation - such as how morphogen signaling regulates the activity of cellular growth pathways, or systemic mechanisms of growth regulation — such as how endocrine growth factors control growth across the whole animal in response to a changing environment.

These tissue-autonomous and systemic mechanisms of growth regulation do not fully explain the allometric relationships observed during development. Transplantation experiments (4) and growth perturbation experiments in *Drosophila* and other insects (1, 5, 6) provide evidence that inter-organ communication may be another mechanism facilitating allometric growth regulation. Based on these observations, Stern and Emlen (7) proposed a model for growth coordination that operates through communication between growing organs, either directly or through a centralized translator like an endocrine organ. However, the mechanisms of this communication pathway have remained unknown.

In *Drosophila* larvae, the growth of the imaginal discs is tightly regulated in order to produce adult structures with specific size and relative proportion (8, 9). This maintenance of allometry is preserved even when developmental growth programs are altered. For instance, *Drosophila* imaginal discs have a remarkable capacity to regenerate and restore proper size and allometry following damage (10, 11). Damage to an imaginal disc activates a regeneration checkpoint (12) that extends the larval period of development (12-16), allowing time for regenerative tissue repair. Regeneration checkpoint activation also slows the growth rate of undamaged tissues (4, 5, 7, 17), coordinating the additional growth and time required for regeneration with the growth of undamaged imaginal discs. Developmental delay and growth coordination are both mediated by expression of *Drosophila* insulin-like peptide 8 (Dilp8) in damaged tissues (18, 19). Dilp8 is a secreted protein that shares structural features with the insulin/relaxin protein family. Several questions remain about how Dilp8 produces both growth regulation and developmental delay. It is possible that these two responses might be mechanistically linked - for instance, growth restriction may lead to developmental delay (14, 20). Alternatively, these two systemic responses may reflect distinct Dilp8-dependent mechanisms. Additionally, it remains unclear whether Dilp8 functions to directly coordinate growth between tissues, or whether Dilp8 mediates growth coordination indirectly through other systemic growth signals.

Nitric oxide synthase (NOS) produces nitric oxide (NO), a potent free radical functioning in all animals to regulate multiple biological processes, including neuronal activity, immunity, and vascular regulation. Experimentally altering the activity of the sole NOS protein found in *Drosophila* produces changes in imaginal disc growth (21) and larval tissue growth (22). However, the mechanism of this regulation remains unknown.

In the experiments presented here, we outline a pathway through which tissues communicate with each other to produce allometric growth. We demonstrate that NOS activity is required for the Dilp8-dependent coordination of growth between regenerating and undamaged tissues following tissue damage, and that NOS regulates growth in undamaged tissues by reducing ecdysone biosynthesis in the prothoracic gland.

## RESULTS

### Nitric oxide synthase activity in the prothoracic gland regulates imaginai disc growth

NOS has been demonstrated to be a regulator of imaginal disc growth during *Drosophila* development (21), but the mechanism of this regulation is unknown. To determine the role of *NOS* and nitric oxide signaling in imaginal disc growth regulation, we systemically misexpressed *NOS* as a transient pulse early in the third larval instar (80hrs after egg deposition [AED]) using Gal4-UAS expression driven by the heat shock promoter (*hs>NOS*). Consistent with previous observations demonstrating that NOS activity restricts imaginal disc growth at the larval to pupal transition (21), we find that transient, systemic misexpression of NOS also reduces imaginal tissue growth during the larval third instar (Fig. 1A, Fig. 1 - supplement 1A). However, targeted overexpression of NOS within imaginal disc tissues produces no observable effect on imaginal disc growth during the third larval instar (Fig. 1 - supplement 2), suggesting that NOS regulates growth via a non-autonomous pathway. Additionally, systemic NOS induction following heat shock produces a developmental delay (Fig. 1B, Fig. 1 - supplement 1B) without producing damage or apoptosis within the imaginal discs (Fig. 1 - supplement 3).

**Figure 1.**
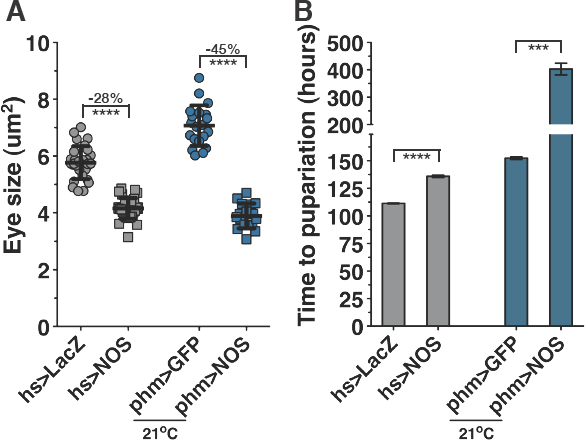
*NOS* overexpression in the prothoraeie gland (PG) regulates imaginai disc growth and developmental timing. (A) *NOS* overexpression restricts imaginai disc growth. Measurement of eye imaginai disc sizes following systemic (*hs>NOS*) or PG-targeted (*phm>NOS*) expression of *NOS.* Heat shock was performed at 76hrs AED (after egg deposition). The area of eye imaginai discs was measured at 104hrs AED for *hs-GAL4* experiments, and 142hrs AED for *phm-GAL4* experiments. *phm>GFP* and *phm>NOS* expressing larvae were raised at 21°C. (B) *NOS* overexpression in the PG promotes developmental delay. Measurement of pupariation timing (marked by eversion of anterior spiracles) following systemic *(hs>NOS)* or PG directed (*phm>NOS*) expression of *NOS.* Statistical analysis: A, mean +/- SD. B, mean of triplicate experiments +/- SEM. ^***^P<0.005, ^****^P<0.001 calculated by two-tailed Student’s t-test.

The timing of developmental transitions in *Drosophila* larvae is regulated by the prothoracic gland (PG) through production of pulses of the steroid hormone ecdysone (23). NOS expression in the PG has been demonstrated to promote the larval to pupal transition by promoting ecdysone production in postfeeding larvae (22). However, when Cáceres et al. constitutively overexpressed NOS in the PG at 25°C throughout larval development, they observed an inhibition of developmental progression and a decrease in larval size. We therefore wondered if NOS might regulate growth through activity in the PG during the larval growth period prior to the post-feeding phase of larval development. Using the *phantom-Gal4* driver, which specifically targets Gal4-mediated expression to the PG throughout larval development (24), we observed that most *phm>NOS* larvae raised at 25°C died prior to the third larval instar. To determine if this larval lethality could be reduced by limiting the expression of NOS in the PG, we raised phm>NOS larvae and control larvae at 21°C. We observed that the majority of *phm>NOS* larvae raised at 21°C progressed through the 3rd instar to pupation (Fig. 1 - supplement 4). We examined the effect on growth when NOS was expressed in the PG of *phm>NOS* larvae and observed that the growth rate of the eye imaginal tissues is reduced relative to control larvae also raised at 21°C (Fig. 1A). These larvae also exhibit a developmental delay when compared to control larvae (Fig. 1B) similar to what is observed during transient, systemic *NOS* misexpression (*hs>NOS*). Therefore, we conclude that *NOS* overexpression in the PG is sufficient to reduce the growth of imaginal discs during the third larval instar and can delay the timing of exit from larval development.

### NOS activity in the PG is necessary to regulate growth during the regeneration checkpoint

During larval development, imaginal disc damage activates a regeneration checkpoint that coordinates the regenerative repair of damaged imaginal tissues with developmental progression. Activation of the regeneration checkpoint produces both 1) a delay in the timing of the larval-pupal transition (12-16), and 2) a reduced growth rate of undamaged imaginal tissues (5, 7, 17). We observe both of these responses upon overexpression of NOS in the PG, without any evidence of imaginal tissue damage. Therefore, we hypothesized that NOS might function during the regeneration checkpoint to mediate developmental delay and growth inhibition in response to imaginal tissue damage.

NOS catalyzes the production of the free radical nitric oxide (NO), an important cellular signaling molecule, from L-arginine. To determine whether NOS activity is increased in the PG during the regeneration checkpoint, we used the fluorescent reporter molecule 4,5-diaminofluorescein diacetate (DAF2-DA) to measure NO production (Fig. 2 - supplement 1). Using this reporter, we observed that larva with genetically targeted wing ablation (*Bx>eiger*), produce increased levels of NO signaling in the PG when compared to undamaged control larvae (*Bx>LacZ*) (Fig. 2A), suggesting that the regeneration checkpoint increases NOS signaling in the PG. To examine whether this activation of NOS in the PG is required for the regeneration checkpoint phenotypes, we expressed a NOS- targeted RNAi to disrupt NOS function specifically in the PG using the *phm-GAL4* driver (*phm>NOS*^*IR-X*^ (22) *orphm>NOS*^*Ri*^ [Bloomington #28792]). To determine the effect of this NOS knockdown in the PG on both the growth of undamaged tissues and developmental timing following X-irradiation, we developed a new technique. Using lead tape to protect the anterior tissues (including the eye-antennal discs) from irradiation-induced damage, we were able to target irradiation damage to the posterior tissues of the larvae. We then visualized the effects of posterior damage on the growth of the shielded and undamaged eye discs located at the anterior end of the developing larvae (Fig. 2B and methods for a description of this technique). Using this shielding technique, we observed that depletion of *NOS* in the PG by RNAi restored eye imaginal disc growth in shielded larvae to the rate observed in unirradiated larvae (Fig. 2C). We observed similar results in larvae homozygous for a loss-of-function allele of NOS (*NOS*^1^ (25)) (Fig. 2 - supplement 2). This demonstrates that NOS function in the PG is necessary to regulate imaginal tissue growth during the regeneration checkpoint.

**Figure 2.**
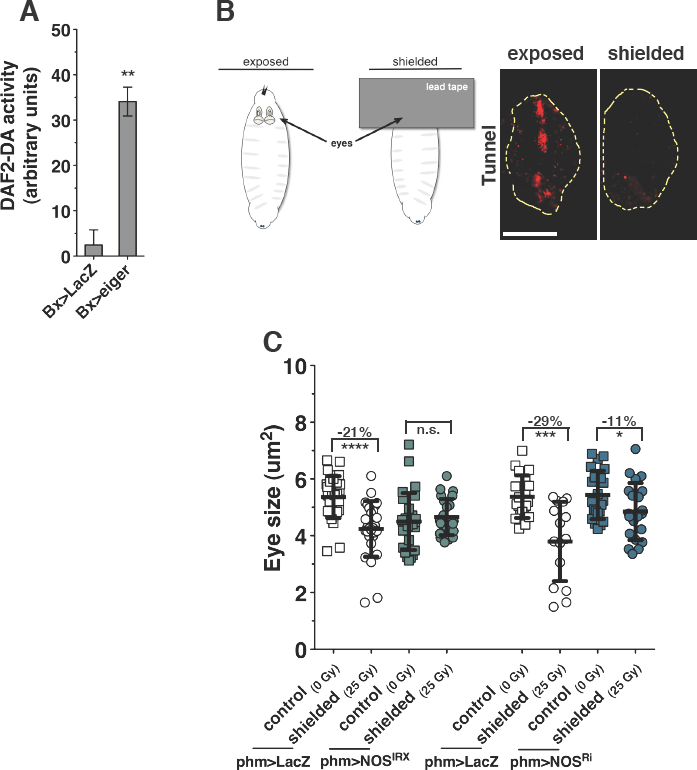
NOS is required in the PG to coordinate imaginai tissue growth during the regeneration checkpoint. (A) Targeted tissue damage (*Bx>eiger*) increases NO production in the PG. Measurement of nitric oxide (NO) production by the fluorescent reporter DAF2-DA. Brain complexes with the PG (outline) were isolated and stained with DAPI and DAF2-DA at 93hrs AED. (B) Illustration of the shielded irradiation method that produces damage in the posterior tissues while protecting anterior tissues from ionizing radiation. Lead shielding protects eye imaginai discs from X-irradiation induced apoptosis. Levels of apoptosis measured by TUNEL staining (red) in eye imaginai discs (outline) isolated from larvae either completely exposed to X-rays, or partially shielded with lead tape to protect anterior tissues from direct damage. (C) *NOS* is required in the PG for regeneration checkpoint growth inhibition. Measurement of undamaged eye imaginai disc size following shielded irradiation (25 Gy) compared to unirradiated control (0 Gy). Posterior tissues were exposed to 25 Gy ionizing irradiation at 72hrs AED and anterior tissues, including the eye discs, were shielded using lead tape. Statistical analysis: A, mean of triplicate +/- SEM. C, mean +/- SD. ^*^ p<0.05, ^**^ p<0.01, ^****^P<0.001 calculated by two-tailed Student’s t-test. Scale bars = 100μm

Both genetic and irradiation damage models depend on the expression of the secreted peptide Dilp8 from damaged tissues for activating the growth restriction and delay of the regeneration checkpoint (Fig. 3 - supplement 1). Additionally, systemic misexpression of *dilp8* is sufficient to induce restriction of imaginal disc growth and developmental delay (Fig. 3 - supplement 2). To examine whether Dilp8 is responsible for activating NOS signaling in the PG, we measured NO production in *Tub>dilp8* misexpressing larvae with the DAF2-DA assay. We observe that *dilp8* misexpression in the absence of damage is sufficient to increase NO production in the PG (Fig. 3A). To determine directly whether *dilp8* induced growth restriction is dependent on NOS function, we measured growth restriction in *NOS* mutant larvae during *dilp8* overexpression in the wing imaginal discs (*Bx>dilp8;NOS*^1%^). We observe that Dilp8 induced growth restriction is reduced in larvae mutant for NOS (Fig. 3B). These results demonstrate that Dilp8 is sufficient to activate NOS in the PG, and that Dilp8 is dependent on *NOS* for imaginal disc growth restriction.

**Figure 3.**
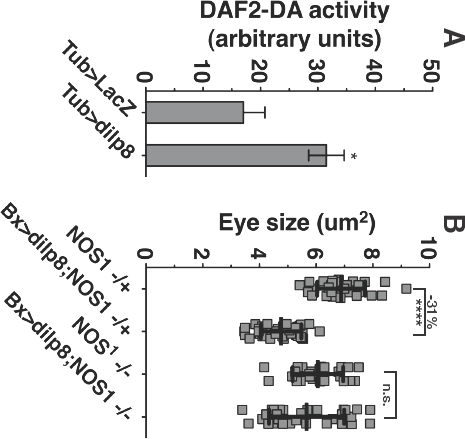
Dilp8 requires NOS to coordinate imaginai tissue growth during the regeneration checkpoint. (A) Systemic *dilp8* expression (*Tub>dilp8*) increases NO production in the PG. Measurement of nitric oxide (NO) production by the fluorescent reporter DAF2-DA. Brain complexes with the PG (outlined) were isolated and stained at 93hrs AED. (B) Imaginai disc growth restriction during wing disc *dilp8* expression (*Bx>dilp8*) is reduced in larvae mutant for *NOS* (*NOS*^1%^). Larvae were raised at 29°C and dissected at lOOhrs AED. Statistical analysis: A, mean +/- SEM. B, mean +/- SD. * P<0.05, ^****^P<0.001 calculated by two-tailed Student’s t-test. Scale bar = 100μm

While growth regulation during the regeneration checkpoint is dependent on NOS function, we observed no effect on the delay of development induced by irradiation in the *NOS* knockdown or in the *NOS* mutant (Fig. 4). Therefore, while Dilp8-dependent growth coordination between regenerating and undamaged tissues requires NOS activity in the PG, Dilp8-dependent developmental delay is mediated by a distinct mechanism. Together, these data demonstrate that localized tissue damage produces two effects: 1) growth inhibition in undamaged imaginal tissues, which is dependent on NOS function in the PG, and 2) a delay in developmental timing, which occurs through a NOS-independent pathway.

**Figure 4.**
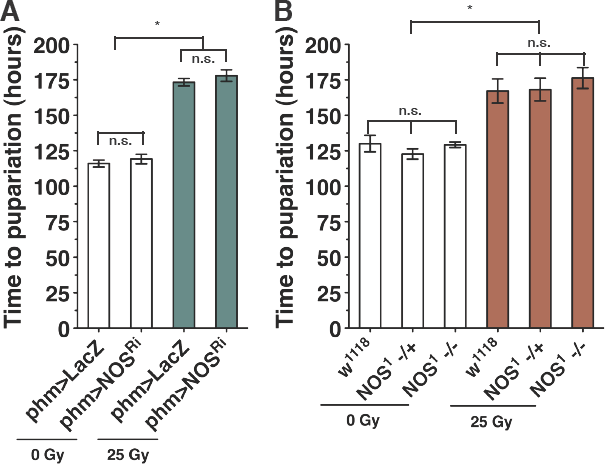
*NOS* is not required for regulation of developmental time during the regeneration checkpoint. LOSS of *NOS* function either by (A) knockdown of *NOS* in the PG (*phm>NOS*^Ri^) or (B) *NOS* mutant (*NOS* ^1^) does not alter developmental delay. Measurement of pupariation timing for larvae with irradiation damage (25 Gy) and control larvae (0 Gy). Mean of triplicates +/- SEM. ^*^ p<0.05 calculated by one-way ANOVA with Tukey’s post-test.

### Imaginal tissues are selectively targeted for growth regulation during the regeneration checkpoint

Larval size is determined by the growth of polyploid larval tissues such as the larval epidermis, fat body, and salivary glands (26). Unlike the diploid imaginai tissues, which become much of the adult fly following metamorphosis, most larval tissues are histolysed during metamorphosis and do not contribute significantly to adult structures. To determine whether imaginal tissues are selectively targeted for growth regulation during the regeneration checkpoint, we compared the effects of checkpoint activation on the growth of imaginal tissues to the effects on the growth of the total larval size, which correlates well with the growth of the polyploid larval epidermis (27). Consistent with our earlier observations, X-irradiation of partially shielded larvae or genetically targeted ablation of wing imaginal discs (*Bx>eiger*) is sufficient to activate the regeneration checkpoint and produce growth inhibition of undamaged eye imaginal discs (Fig. 5A, B). Both damage models depend on the damaged tissues expressing the secreted peptide Dilp8 for growth inhibition and developmental delay (Fig. 3). Consistent with a role for NOS and Dilp8 in growth regulation during the regeneration checkpoint, we observe growth inhibition of undamaged eye imaginal tissues by expression of *NOS* in the PG (*phm>NOS*, Fig. 5B) and *dilp8* expression in the wing pouch, the region of the imaginal disc that will become the adult wing blade (*rn>dilp8*, Fig. 5B).

**Figure 5.**
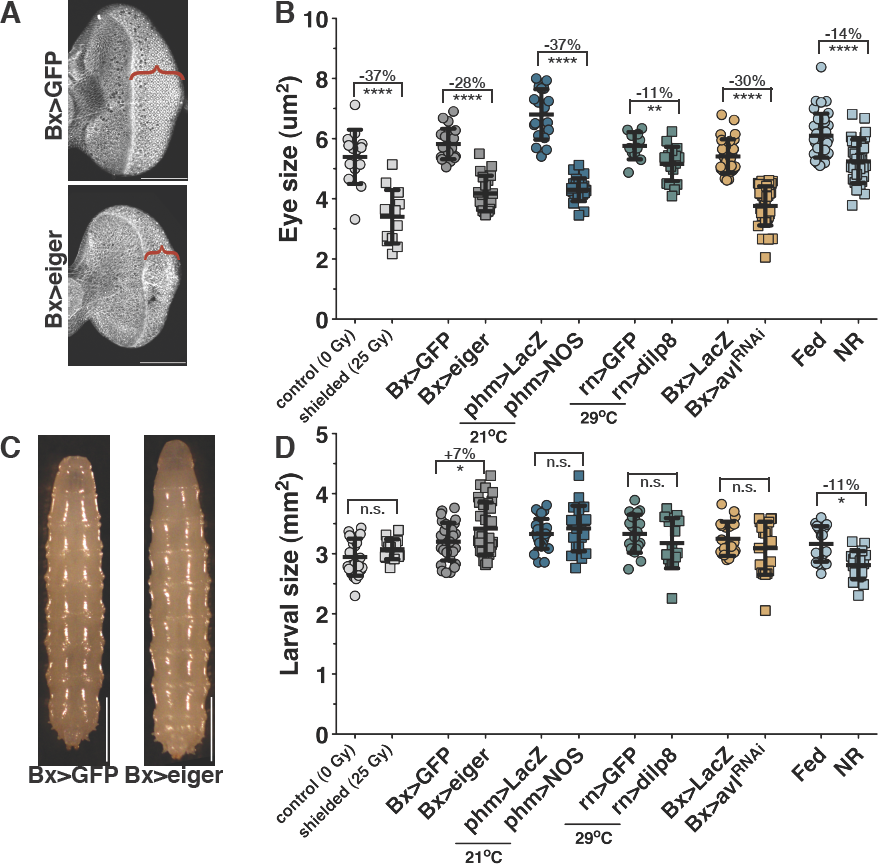
The regeneration checkpoint selectively restricts imaginal tissue growth. (A) Growth reduction and developmental delay of undamaged eye imaginal discs in larvae with targeted tissue damage in the wings (*Bx>eiger*) and control larvae (*Bx>GFP*). Wings were isolated at 104hrs AED and stained with rhodamine-labeled phalloidin. Brackets highlight the position of the morphogenetic furrow in each disc. Scale bar = 100 μm. (B) Measurement of eye imaginal disc size following nutrient restriction (NR) or multiple distinct activators of the regeneration checkpoint including: shielded irradiation with 25 Gy (shielded), expression of pro-inflammatory signal (*Bx>eiger*), expression of NOS in the PG (*phm>NOS*), wing-targeted expression of dilp8 (*rn>dilp8*), and wing-targeted neoplastic transformation (*Bx>avlRNAi*). Larvae were raised at 25°c and eye imaginal discs were isolated at 104hrs AED for measurement for all experiments except for the following: *phm>LacZ* and *phm>NOS* larvae were raised at 21°C to reduce NOS overexpression and permit analysis of third instar growth phenotypes. *rn>GFP* and *rn>dilp8*, raised at 29°C and eye discs were dissected and measured at 80hrs AED to maximize dilp8 overexpression and the systemic growth phenotype. (C) The regeneration checkpoint does not restrict larval growth. *Bx>eiger* and control larvae isolated at 104hrs AED. Scale bar = 1 mm. (D) Measurement of larval growth. Larvae were raised and isolated for measurement as in B. Statistical analysis: B and D, mean +/- SD. ^*^ p<0.05, ^***^p<0.005, ^****^p<0.001 calculated by two-tailed Student’s t-test.

In contrast, we found that checkpoint activation does not reduce overall larval growth (Fig. 5C, D). In our two damage models, larval growth continued at the same rate or even slightly faster than the growth observed in control larvae. Similarly, we observed a slight but not statistically significant increase in the rate of larval tissue growth in larvae with *phm>NOS* and wing-targeted expression of Dilp8 (*rn>dilp8*), as compared with control larvae (Fig. 5D). Additionally, we examined other disruptions of wing imaginal disc growth and found that induction of neoplastic tumors in the wing imaginal tissues using knockdown of the *Drosophila* syntaxin protein Avalanche (*Bx>avi*^*RNAi*^, (28)) also produces slower growth in the eye imaginal discs without altering larval tissue growth. This is similar to what is observed following activation of the regeneration checkpoint (Fig. 5B, D), consistent with the observed activation of Dilp8 during tumorigenesis (18, 19). This pattern of growth regulation observed during the regeneration checkpoint - reduced growth of imaginal discs and sustained or even increased growth of larval tissues - contrasts with the growth pattern observed in larvae experiencing reduced insulin signaling in response to nutrient restriction, where growth of both imaginal and larval tissues are reduced (Fig. 5B, D). Therefore, we sought another growth regulatory pathway that would explain how the regenerative checkpoint could specifically reduce imaginal disc growth.

### The regeneration checkpoint reduces growth of undamaged tissues by limiting ecdysone signaling

The PG produces pulses of ecdysone synthesis during the larval growth phase that determine the timing of developmental transitions such as larval molts, the mid-third instar transition (29), critical weight (30), and the exit from larval development. Experimental evidence exists to support roles for ecdysone in both promoting (5, 31-33) and restricting (24, 32-35) growth of imaginal discs. Since overexpression of NOS in the PG influences both developmental timing and imaginal disc growth, we examined whether NOS activity in the PG alters ecdysone signaling during the regeneration checkpoint. To examine ecdysone signaling during larval development, we measured the expression of the ecdysone transcriptional targets *E74B* and *E74A.* Transcription of *E74B* is induced by low pulses of ecdysone that occur in between molts, during the larval third instar, and inhibited by high levels of ecdysone, whereas *E74A* transcription is induced by high levels of ecdysone such as the large pulse of ecdysone that triggers the exit from larval development (36). Levels of *E74B* transcription are often used as a measure of ecdysone activity during the larval growth phase (5, 34, 37). Consistent with previous studies (37), we observed that transcription of *E74B* is reduced following activation of the regeneration checkpoint in *Bx>eiger* expressing larvae (Fig. 6A, Fig. 6 - supplement 1A). In larvae overexpressing NOS in the PG (*phm>NOS*) we observed that the expression of E74B is lower than in control larvae during the mid 3rd instar (Fig. 6B 140hrs), suggesting that ecdysone titers are reduced. This interpretation was confirmed with a direct measure of ecdysone levels using a competitive enzyme immunoassay (38) which demonstrated that ecdysone levels are lower in *phm>NOS* compared to control larvae of the same age. Additionally, E74A expression is not substantially increased in *phm>NOS* larvae (Fig. 6 - supplement 1B) consistent with the reduced ecdysone levels that we observed in *phm>NOS* larvae. Together, these results suggest that ecdysone signaling is reduced when NOS is active in the PG during the 3rd instar larval growth period.

**Figure 6.**
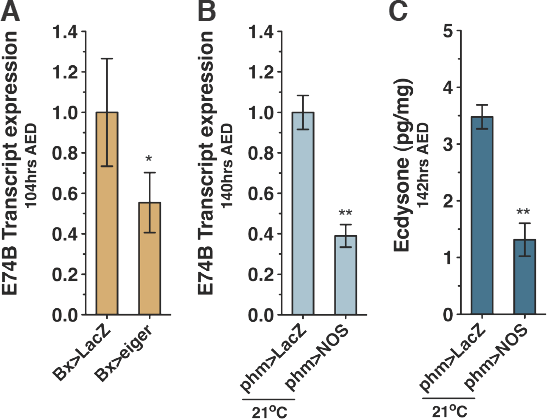
*NOS* overexpression in the PG reduces eodysone signaling and eodysone production. (A) Regeneration checkpoint activation reduces eodysone signaling. Transcription of ecdysone-induced *E74B,* a reporter for early ecdysone levels during the third larval instar, in control larvae (*Bx>LacZ*) and larvae with targeted tissue damage (*Bx>eiger*), (B) *NOS* activity in the PG reduces ecdysone signaling. Transcription of *E74B* is reduced in larvae with *NOS* overexpression in the PG (*phm>NOS*) compared to control (*phm>LacZ*) larvae. Transcription levels measured by qRT-PCR in triplicate, normalized to control expression levels. (C) *NOS* activity in the PG reduces ecdysone production. The presence of ecdysone is reduced in larvae with *NOS* overexpression in the PG (*phm>NOS*) compared to control (*phm>LacZ*) larvae. Ecdysone levels measured by ELISA assay isolation triplicate. Statistical analysis: mean of triplicates +/- SEM. ^*^ p<0.05, ^**^ p<0.01, calculated by paired one-tailed t-test.

We then determined whether the reduced growth of undamaged discs during regeneration checkpoint activation is directly dependent on the reduced ecdysone expression that we observed. Feeding larvae food supplemented with ecdysone (0.6mg/ml) effectively increases ecdysone titer in larvae during the feeding period (34). Using this approach, we tested whether we could bypass NOS-dependent growth inhibition by ecdysone feeding. We observed that ecdysone feeding can bypass the imaginal disc growth restriction produced by: 1) imaginal tissue damage (*Bx>eigei*) 2) regeneration checkpoint signaling (*Tub>dilp8*), or 3) misexpression of *NOS* (*hs>NOS*) and overexpression of *NOS* in the PG (*phm>NOS*) (Fig. 7A and Fig. 7 - supplement 1A). This level of ecdysone feeding did not significantly alter the growth of larval tissues during damage or NOS overexpression, but strongly reduced larval tissue growth in *dilp8* misexpressing larvae, as reflected in the overall larval size (Fig. 7 - supplement 2A).

**Figure 7.**
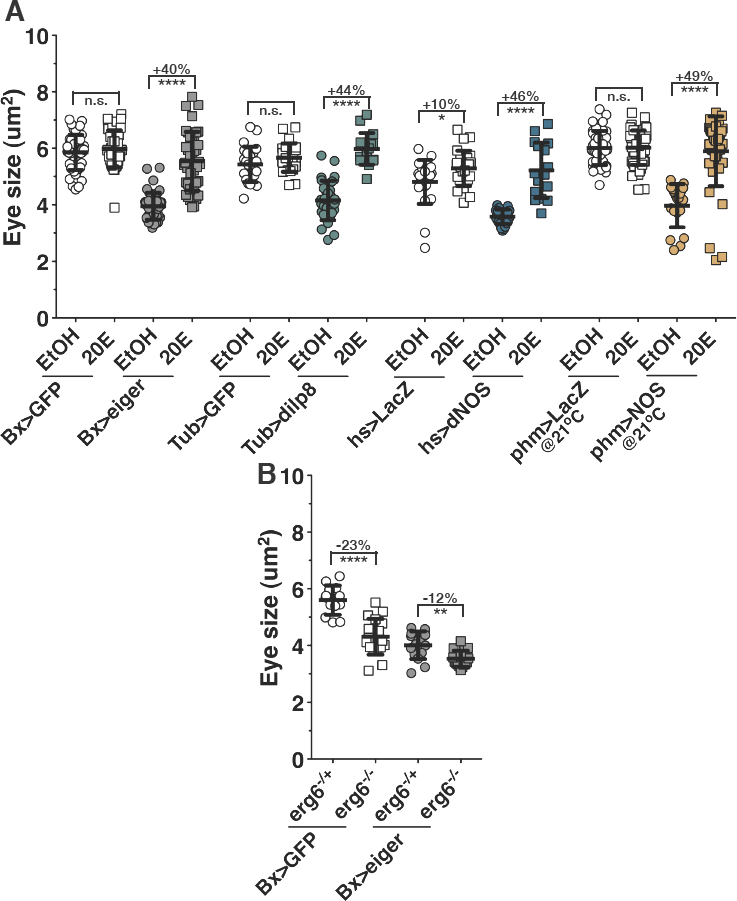
Imaginal disc growth restriction during the regeneration checkpoint is the result of reduced ecdysone signaling. (A) Ecdysone levels are rate-limiting for imaginal disc growth during the regeneration checkpoint. 20-hydroxyecdysone (20E) rescues growth restriction induced by Bx>eiger, systemic *dilp8* misexpression (*Tub>dilp8*), and *phm>NOS* compared to control ethanol only fed larvae (EtOH). (B) Nonadditive effects of wing damage and ecdysone reduction on eye imaginal disc growth suggest convergent mechanisms. Measurement of eye imaginal disc size in larvae with targeted wing damage (*Bx>eiger*) and control larvae (*Bx>GFP*). Larvae were transferred at 80hrs to food lacking steroid ecdysone precursor (*erg6* +/-) or control food (*erg6* +-/). Eye imaginal discs were dissected at 104hrs AED and measured. Statistical analysis: mean +/- SD. ^**^ p<0.01, ^****^p<0.001 calculated by two-tailed Student’s t-test.

To directly determine whether ecdysone promotes imaginal disc growth during normal development, we reduced ecdysone levels in third instar larvae by transferring larvae to yeast-sucrose food prepared using *erg6*^-/-^ mutant yeast, which lacks the necessary steroid precursors for ecdysone synthesis (39, 40) (see Experimental Procedures). This resulted in a marked decrease in imaginal tissue growth (Fig. 7B), an extended developmental time to pupation (Fig. 7 - supplement 2C), and a slight, but significant increase in larval tissue growth (Fig. 7 - supplement 2D), as compared with larvae reared on similar food derived from heterozygous *erg6*^-/+^ yeast. Therefore, ecdysone is required for a normal rate of imaginal disc growth even in the absence of imaginal disc damage. Furthermore, we observed that ecdysone limitation by growth on *erg6*^-/-^ food produces only a minor effect on the growth of imaginal tissues in *Bx>eiger* larvae (Fig. 7B). This epistatic interaction supports a model in which the regeneration checkpoint and ecdysone regulate imaginal tissue growth via convergent mechanisms.

### NOS activity in the PG regulates expression of ecdysone biosynthesis genes

To better understand how NOS activity in the PG reduces ecdysone signaling in larvae, we examined whether NOS regulates the expression of ecdysone biosynthetic genes. Ecdysone is synthesized in the PG from sterol precursors by the consecutive actions of the P450 enzymes collectively referred to as the Halloween enzymes (41). Previous work has demonstrated that the expression of Halloween genes is reduced during activation of the regeneration checkpoint (37). To determine whether NOS regulates ecdysone synthesis by limiting Halloween gene expression, we examined the effect on the transcription of the Halloween genes *spookier* (*spok*) (42) and *disembodied* (*dib*) during either targeted tissue damage or NOS overexpression in the PG (43). Transcription of both *spok* and *dib* are reduced in *Bx>eiger* larvae and *phm>NOS* larvae in comparison to control larvae (Fig. 8A,B) when examined during the larval growth phase, a time when we observe reduced ecdysone signaling following activation of the regenerative checkpoint or NOS misexpression in the PG. Therefore, upon activation of the regeneration checkpoint, NOS functions in the PG to reduce ecdysone signaling through the transcriptional repression of ecdysone biosynthesis genes.

**Figure 8.**
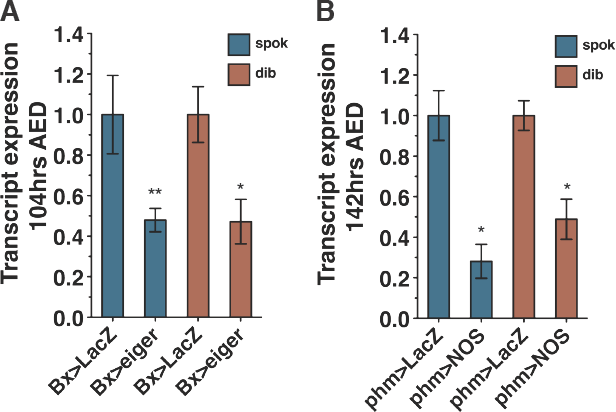
Regeneration checkpoint activation and *NOS* overexpression in the PG both reduce transcription of Halloween genes. (A) Regeneration checkpoint activation reduces Halloween gene transcription. Expression of the Halloween genes spookier (*spok*) and disembodied (*dib*), in control larvae (*Bx>LacZ*) and larvae with targeted tissue damage (*Bx>eiger*). (B) NOS activity in the PG reduces Halloween gene transcription. Expression of *spok* and *dib* in control (*phm>LacZ*) larvae and larvae with *NOS* overexpression in the PG (*phm>NOS*). Larvae were hatched and raised at 21°C. Transcription levels measured by qRT-PCR in triplicate, normalized to control transciption levels. Statistical analysis: A and B mean of triplicates +/- SEM. ^*^ P<0.05, ^**^ P<0.01, calculated by paired one-tailed student’s t-test.

### NOS inhibition of ecdysone synthesis is limited to the feeding phase of larval development

Based on our observations, we conclude that during the growth phase of larval development, NOS activity in the PG restricts the expression of ecdysone biosynthesis genes, reducing ecdysone synthesis. However, this model contrasts with a model arising from previous work demonstrating that NOS activity in the PG of post-feeding larvae enhances ecdysone production by inhibiting the nuclear hormone receptor *E75,* an antagonist of ecdysone biosynthesis (22). To reconcile these two distinct descriptions of NOS activity in the PG, we first sought to determine whether this post-feeding pathway could be active during the growth phase of larval development. We observed that the *E75B* expression, which is normally upregulated during the post-feeding period of larval development, is completely suppressed in larvae with targeted wing damage (Fig. 9 - supplement 1). Therefore, we conclude the pathway by which NOS can promote ecdysone production in post-feeding larvae is not likely to be active during the growth phase of larval development and is delayed following activation of the regeneration checkpoint. Consistent with this idea, we observed that the ability of *NOS* misexpression to delay pupation was most robust when expressed during larval feeding (76hrs or 80hrs). This delay was significantly decreased when *NOS* was misexpressed later in the 3rd instar as the larvae entered the postfeeding phase (96hrs or 104hrs) (Fig. 9A - supplement 2A). This suggests that the ecdysone-inhibiting and ecdysone-promoting mechanisms of NOS might be temporally separated during the larval growth and post-feeding phases of development. To test this hypothesis, we determined whether the reduction of imaginal disc growth and delay of developmental time induced by transient systemic misexpression of *NOS* (*hs>NOS*) are dependent on the phase of development. We observed that misexpressing *NOS* early in the 3rd instar during the larval feeding period (76hrs AED) produces a robust developmental delay and restriction of imaginal disc growth (Fig. 9B and B - 76hrs, Fig. 9 - supplement 2B). However, we found that misexpression of *NOS* late in the 3rd instar, at the time that larvae stop feeding (104hrs AED), produces minimal effect on developmental time and imaginal discs (Fig. 9A and B - 104hrs), with a slight increase in growth, which would be consistent with a slight increase in ecdysone signaling. This led us to test whether a pulse of NOS early in development would inhibit ecdysone biosynthesis as our model would predict and whether a pulse late in development would increase ecdysone biosynthesis late in development, as the Cáceres model predicts. We measured ecdysone signaling and Halloween gene transcription after an early or late heat shock (76hrs vs 104hrs). We observed that the 76hrs pulse of *NOS* decreased *E74B* transcription suggesting that ecdysone signaling is inhibited. We observed that the 76hr pulse of *NOS* inhibited *dib* but not *spok* transcription. When we induced a late pulse of *NOS* (104hrs), we observed that *E74B* and *dib* were not decreased, while *spok* transcription increased (Fig. 9C and D). These data are consistent with our growth and developmental time results, as well as the previously published Cáceres model. These results suggest that as development progresses, the regulatory effect of NOS in the PG has two distinct states. During the feeding phase of larval development, NOS inhibits ecdysone production as we describe here (Fig. 10). Later, in post-feeding larvae, NOS functions to promote ecdysone synthesis by inhibiting E75 activity as described in previous work (22).

**Figure 9.**
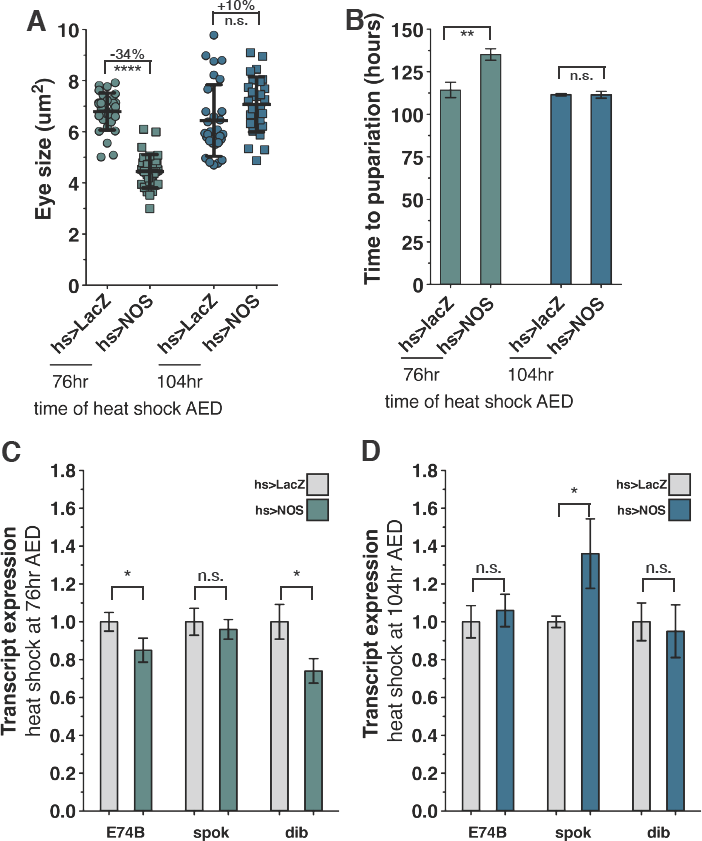
NOS activity during the larval feeding period inhibits ecdysone and *NOS* activity during the larval wandering period promotes ecdysone. (A) Expression of *NOS* early during the larval feeding period restricts imaginal disc growth, while *NOS* expression late during the wandering period does not inhibit growth. *NOS* was systemically expressed by heat shock (*hs>NOS*) at either 76hrs or 104hrs AED and eye imaginal disc size was measured at 116hrs AED. (B) Expression of *NOS* early during the larval feeding period delays larval development, while *NOS* expression late during the wandering period does not delay development. *NOS* was systemically expressed by heat shock (*hs>NOS*) at either 76hrs or 104hrs AED and time to pupation was measured. (C) Expression of *NOS* early during the larval feeding period restricts ecdysone biosynthesis genes and signaling. *NOS* was systemically expressed by heat shock (*hs>NOS*) at 76hrs AED and ecdysone signaling (E74B) and Halloween gene (*spok* and *dib*) transcription was measured by qRTPCR at 116hrs AED. (D) Expression of NOS late during the wandering period promotes ecdysone biosynthesis gene transcription. *NOS* was systemically expressed by heat shock (*hs>NOS*) at 104hrs AED and ecdysone signaling (E74B) and Halloween gene (*spok* and *dib*) transcription was measured by qRT-PCR at 116hrs AED. Statistical analysis: A, mean +/- SD. B, mean of triplicates +/- SEM. ^*^ p<0.05, ^**^ p<0.01, ^***^p<0.0005, ^****^p<0.001 calculated by two-tailed Student’s t-test. C and D mean of duplicates +/- SEM. ^*^ p<0.05 calculated by paired one-tailed student’s t-test.

**Figure 10.**
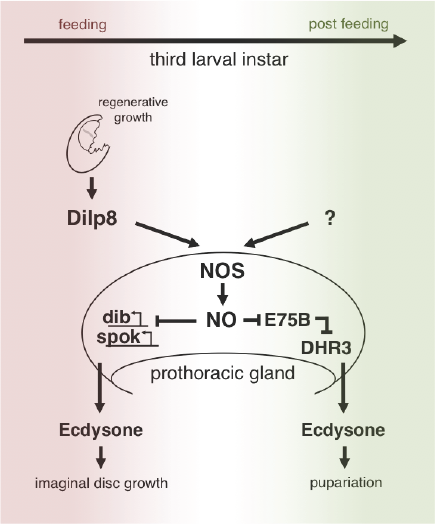
Model for allometric growth regulation by NOS.

## DISCUSSION

During *Drosophila* development, damage to larval imaginal discs elicits a regeneration checkpoint that has two effects: 1) it delays the exit from the larval phase in development to extend the regenerative period, and 2) it coordinates regenerative growth with the growth of undamaged tissues by slowing the growth rate of distal, undamaged tissues. How the damaged and regenerating tissues communicate with undamaged tissues to coordinate growth has been an open question. Perhaps damaged tissues produce signals that directly influence the growth rate of undamaged tissues. Alternatively, damaged tissues may indirectly influence the growth of undamaged tissues by producing signals that alter the systemic expression of limiting growth factors. Consistent with the latter possibility, we describe a novel indirect-communication pathway for growth coordination that functions during the regeneration checkpoint (Fig. 10).

An essential component of this growth coordination is the secreted peptide Dilp8, which is released by damaged tissues and is both necessary and sufficient to regulate the growth of distal tissues during the regeneration checkpoint (18, 19). Dilp8 shares structural similarity to insulin-like peptides, which function to stimulate growth by activating the insulin receptor. However, in contrast to insulin-like peptides, Dilp8 acts to limit growth. A simple model explaining Dilp8 function would be that Dilp8 acts directly as an antagonist to insulin receptor activity, thus reducing growth in undamaged tissues. However, we show that the growth response to checkpoint activation of polyploid larval tissues differs from imaginal discs (Fig. 5). These results are inconsistent with Dilp8 regulating imaginal disc growth through antagonizing systemic insulin signaling.

In contrast, we show here that NOS functions in the PG to regulate the growth of imaginal discs in response to Dilp8 expression from damaged tissues during the developmental checkpoint. We demonstrate that growth coordination during the regeneration checkpoint depends on *NOS* gene function in the PG. We show that damage, as well as *dilp8* expression on its own, is able to increase NO production in the PG. NOS activity inhibits the growth of undamaged imaginal discs by reducing ecdysone synthesis in the PG. We show that NOS does this through the transcriptional inhibition of the P450 enzymes *disembodied* and *spookier,* which are necessary for ecdysone biosynthesis. Although it has been known for some time that NOS activity is capable of regulating growth of imaginal discs (21), the experiments here elucidate the mechanism of this growth regulation.

The mechanism we describe here seems to contrast published experiments demonstrating that NO signaling in the PG promotes ecdysone synthesis following the larval feeding period during the larval-to-pupal transition (22). However, experiments from this published work also demonstrate that earlier *NOS* expression in the PG during larval development produces small larvae that arrest at second larval instar stage of development. This arrest can be partially rescued by either ecdysone feeding (22), or by reducing the level of GAL4-UAS driven *NOS* expression by raising larvae at a lower temperature (Fig. 1 - supplement 2). Additionally, other experiments have shown that pharmacological increase of NO levels in larvae can produce larval developmental delays (44, 45). Together, these observations suggest that NOS activity earlier in larval development might inhibit rather than promote ecdysone signaling during the larval growth period. Additionally, we observe that *E75B* is not expressed in larvae with targeted wing ablation (Fig. 6 - supplement 1), suggesting that the pathway for increasing ecdysone may not be available during the regeneration checkpoint. We have focused on the role of NOS during the middle of the third larval instar (76-104h AED) and have found that NOS activity during this period of development inhibits growth and ecdysone, while NOS activity at the end of larval development does not inhibit growth. Based on our results, we conclude that there are distinct roles for NOS in the PG during different phases in development; NOS activity post-larval feeding promotes ecdysone synthesis through inhibition of *E75,* whereas NOS activity during the larval growth phase reduces ecdysone synthesis and signaling. These observations raise interesting questions regarding what signals promote NO production during the larval-to-pupal transition.

Furthermore, we demonstrate here that ecdysone is a rate-limiting promoter of imaginal disc growth during both normal development and during the regeneration checkpoint. Most studies have supported a model in which ecdysone acts as negative regulator of growth based on two observations: 1) the final pulse of ecdysone at the end of the third larval instar shortens developmental time and therefore reduces final organ size, and 2) increased ecdysone signaling can antagonize Dilp synthesis in the fat body. However, when measuring the effects of ecdysone on growth, many previous studies have focused on measuring either the growth of the larvae (which as we observe does not always reflect the growth of the imaginal tissues) or measuring the final size of adults (which is a function of both growth rate and time). When one either examines clones expressing mutant alleles of ecdysone receptor (46) or measures the growth of entire imaginal discs directly following ecdysone feeding as we have done here, ecdysone signaling can be shown to promote imaginal disc growth. Moreover, we have shown that during the regeneration checkpoint, reduced ecdysone levels resulting from NOS activity in the PG are necessary and sufficient to limit the growth of undamaged imaginal discs. Finally, we have observed that damage-induced activation of the regeneration checkpoint slows the progression of the morphogenetic furrow in undamaged eye discs (see Fig. 5A). Since the progression of the furrow is dependent on ecdysone (47), its slow progression is also consistent with the regeneration checkpoint producing reduced ecdysone signaling.

During the regeneration checkpoint, there are two well-characterized responses to imaginal tissue damage, 1) the reduction of undamaged imaginal tissue growth and 2) a delay in the timing of pupation. As described above, the reduction of imaginal tissue growth is dependent on reduced ecdysone signaling during the third larval instar. Since growth coordination and the delay in developmental timing are both dependent on reduced ecdysone levels, we might expect both effects to be dependent on the same mechanism. However, we clearly demonstrate that the genetic requirements for these two systemic responses to damage are distinct. NOS is necessary for growth regulation following tissue damage, but is not necessary for the developmental delay, while Dilp8 is necessary for both growth regulation and developmental delay. While we do observe that overexpression of NOS in the PG produces developmental delay, we interpret these results as showing that high levels of expression of NOS in the PG are capable of suppressing ecdysone synthesis, thus producing delay. Therefore, we conclude that Dilp8 secretion from damaged imaginal discs, which is necessary for both developmental delay and growth coordination, must produce these two responses through distinct mechanisms.

In addition, these observations raise an important question about the regulation of regenerative growth. Since growth coordination between regenerating and undamaged tissues is achieved through reduced ecdysone signaling, which is necessary for imaginal disc growth, then regenerative growth, which is able to proceed despite reduced ecdysone signaling (48), must have distinct growth requirements from undamaged tissues. Understanding these differences in growth regulation could provide valuable insights into the mechanistic distinctions between regenerative and developmental growth.

## MATERIALS AND METHODS

### Drosophila Stocks

*w*^*^; *P{UAS-Nos.L}2; P{UAS-Nos.L}3* was provided by Pat O’Farrell (49). *y,w; phm-GAL4[51A2]* was provided by Alexander Shingleton (24). UAS-NOS^mac^ and *UAS-NOS*^*IR-X*^ was provided by Henry Krause (22). *NOS*^1^ was provided by James Skeath (25). *UAS-eiger* and *UAS-reaper* and *rn-Gal4, UAS-YFP*were provided by Iswar Hariharan (50). *UAS-dilp8::3xFLAG* was provided by Maria Dominguez (19). *UAS-Avl*^*RNAi*^ was provided by David Bilder (28). All other stocks were obtained from the Bloomington *Drosophila* Stock Center or the Vienna *Drosophila* RNAi Center. Identifying stock numbers are referenced in the text.

### Drosophila culture and media

Unless otherwise specified, larvae were reared at 25°C on standard cornmeal-yeast-molasses media (Bloomington *Drosophila* Stock Center) supplemented with live bakers yeast granules after developmental synchronization by egg staging. Developmental timing was synchronized through the collection of a 4 hour egg laying interval on grape agar plates. 20 hatched first-instar larvae were transferred to vials containing media 24hrs after egg deposition (AED) (48hrs AED when raised at 21°C). Heat shock-mediated expression was induced at 76hrs AED by 29°C pretreatment and heat shock for 30min at 37°C. Nutrient restriction was initiated at 92hrs AED by transferring larvae to 1% agarose/PBS for the rest of larval development. Exogenous application of ecdysteroid was preformed as previously described (12). Briefly, larvae were transferred at 80hrs AED (*Bx>eiger, Tub>dilp8, Bx>dilp8*) or 124hrs AED (*phm>NOS*) to either 0.6 mg 20-hydroxyecdysone (Sigma) dissolved in 90% ethanol/ml of media, or an equivalent volume of ethanol alone. For ecdysone restriction assays, a defined yeast media was prepared with the *erg-6* mutant yeast strain, sucrose, and agar (39, 40), and larvae were transferred from standard media to *erg6*^-/-^ or *erg6*^+%^ media at 80hrs AED.

### Ionizing irradiation damage

Irradiation was performed as previously described (12). Briefly, 4 hour staged larvae were raised in petri dishes on standard media and exposed to 25 Gy X-irradiation generated from a Faxitron RX-650 operating at 130kV and 5.0mA. For partially shielded irradiation exposure experiments, shielded and control larvae were immobilized by being chilled in an ice bath, mounted on chilled glass cover slips, and kept on ice during the duration of the irradiation. Larvae were partially shielded from ionizing irradiation by placing a 0.5 cm^2^ strip of lead tape (Gamma) over the estimated anterior third of their body, covering segments T1-T3. Larvae and control larvae were returned to cornmeal-molasses food at 25°C following irradiation.

### Measurement of growth parameters

Time to pupation, the time at which half the population had pupated, was calculated by recording the number of pupariated individuals every 12hrs. For measuring imaginal tissue area, tissues were dissected in phospho-buffered saline (PBS), fixed in 4% paraformaldehyde, mounted in glycerol, imaged by DIC on a Zeiss Axioplan2 microscope, and measured in ImageJ (NIH). The area of staged larvae was imaged on an Olympus DP21 microscope digital camera when viewed from the dorsal aspect and measured in ImageJ.

### Indirect immunofluorescence

Dissected tissues were fixed for 20 minutes in 4% paraformaldehyde, washed in PBS with 0.3% Triton-X100 to permeablize cells, treated with primary antibodies (overnight at 4 °C; rabbit anti-cleaved Caspase-3 (Asp175) 1:100, Cell Signaling Technology, MA), and secondary antibodies (4 hrs at room temperature). Cell death detection by TUNNEL with TMR red fluorescent probe (Hoffmann-La Roche, Basel, Switzerland) was preformed following manufacturer instructions. Briefly, labeling buffers were mixed with secondary antibody stain and incubated for 2hrs at 37°C.

### Ecdysone measurements

Ecdysone levels in third instar larvae were quantified using a competitive enzyme immunoassay (Cayman Chemicals) as described previously (37).

### NADPH-diaphorase assay

NOS activity was detected by measuring NADPH-diaphorase activity through an adapted method (51). Tissues were fixed for 1 hr in 4% paraformaldehyde and then permeablized in 0.3% Triton X-100 for 20min. Fixed tissues were suspended in NADPH-diaphorase staining solution in the dark for 15min, then washed in PBS, mounted in 80% glycerol, and imaged by DIC.

### DAF2-DA assay

NO production was detected by 4,5-Diaminofluorescein diacetate (DAF2-DA, Sigma). Brain complexes were dissected in PBS and incubated in 10uM DAF2- DA for 1 hr at 28°C, rinsed in PBS, stained with DAPI 1: 1000, rinsed in PBS, and imaged with by confocal microscopy. DAF2-DA fluorescence was quantified in ImageJ by measuring the mean gray value of each PG lobe normalized to the background fluorescence of the adjacent brain hemisphere.

### PCR

#### Semi-quantitative PCR

RNA was isolated from staged larvae using TRIzol reagent treatment (Invitrogen-Life Technologies, CA) followed by RNeasy cleanup (Qiagen, Limburg, Netherlands) and DNase treatment with the Turbo DNase-kit (Ambion-Life Technologies, CA). RNA yield was quantified by using UV spectroscopy to measure A260. cDNA template for RT-PCR was generated using 1 μg sample RNA as a substrate for Roche Transcriptor first strand cDNA synthesis using poly dT primers. Polymerase chain reaction (PCR) was performed with TaKaRa Ex Taq DNA Polymerase (Takara, Otsu, Japan) in a MJ research PTC-200 DNA Engine Cycler. Conditions for amplification were as follows: 94°C for 2 minutes, then 94°C for 15 seconds, 50°C for 15 seconds, and 72°C for 15 seconds for 23 cycles with Tubulin primers or 31 cycles with *E75B* primers. Amplified products were then identified by electrophoresis on a 3% agarose gel and visualized with SYBR Green (Life Technologies, CA) through epifluorescent analyzer (Fujifilm Intelligent Lightbox LAS-3000). Relative expression differences were measured in ImageJ in relation to *tubulin* expression. Primers: *E75B* (52), *tubulin* (tub-L CTCATAGCCGGCAGTTCG)(tub-R GAT AGAGAT ACATT CACGCAT ATT GAG).

#### Quantitative RT-PCR

RNA was isolated and cDNA was generated as described above except for Fig. 9 which used ReliaPrep™ RNA Cell and Tissue Miniprep Systems (Promega) and poly dT primers with random hexamer primers. cDNA was analyzed using a Mastercycle EP Replex real-time PCR system (Eppendorf). Fold change was calculated relative to *tubulin* expression by the -ΔΔCt method (53). Isolates were taken from at least three stagings to calculate the mean fold change. Two to three independent RNA isolations were assayed within each staging and used to calculate standard error of the mean across stagings. Primers: *E74B* (18), *spookier* (spo-L CGGTGATCGAAACAACTCACTGG, spo-R GGATGATTCCCGAGGAGAGCAG), *disembodied* (dib-L AGGCTGCTGCGTGAATACG, dib-R TCGATCAGCACTGGAGCATC).

## ACKNOWLEDGEMENTS

We would like to thank I. Hariharan, D. Bilder, M. Dominguez, A. Shingleton, P. O’Farrell, H. Krause, and J. Skeath for *Drosophila* lines, Q. Ou for advice on the DAF2 assays, D. Castle, D. DeSimone, P. Adler, J. Hirsh, D.G. Halme, and I. Hariharan for their suggestions on the manuscript, and S. Bekiranov for advice on statistical analyses.

---

**Figure 1 - supplement 1.**
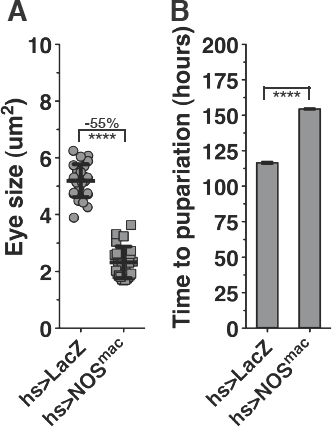
Systemic misexpression of mouse macrophage *NOS* {*NOS*^*mac*^) regulates imaginai disc growth and developmental timing. (A) Misexpression of the mouse-macrophage form of *NOS* (*NOS*^mac^) restricts growth. Measurement of eye imaginal disc sizes in larva with systemic misexpression of *NOS*^mac^ expression (*hs>NOS*^mac^) at 104hrs AED. (B) *NOS*^mac^ misexpression delays development. Statistical analysis: A, mean +/- SD. B, mean of triplicate experiments +/- SEM. ^****^P<0.001 calculated by two-tailed Student’s t-test.

**Figure 1 - supplement 2.**
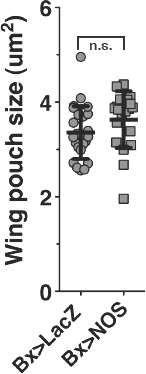
*NOS* overexpression in the wing disc does not reduce growth. Targeted misexpression of *NOS* to the pouch of the wing imaginai tissue (*BX>NOS*) is not sufficient to reduce growth of the wing pouch, nor wing area (data not shown). Wing imaginal discs measured at 104hrs AED from larvae with targeted expression of *NOS* in the wing (*BX>NOS*) and control (*Bx>LacZ*) larvae. Statistical analysis; mean +/- SD. n.s. calculated by two-tailed Student’s t-test.

**Figure 1 - supplement 3.**
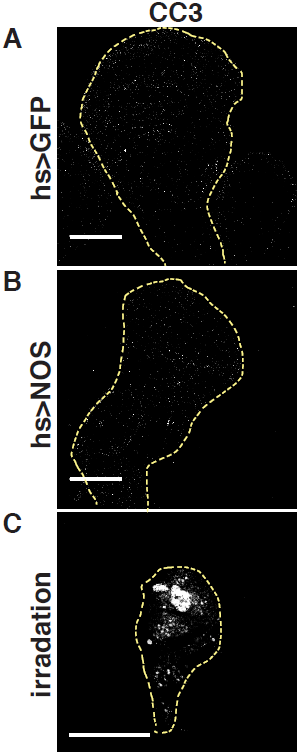
Systemic *NOS* misexpression does not induce cell death. Systemic *NOS* misexpression (*hs>NOS*) does not induce cell death in the wing discs. Cleaved caspase staining (COS) in wing discs (outlines) isolated at 104hrs AED. Control (*hs>GFP*) and *NOS* misexpression larvae (*hs>NOS*) that had been heat-shock treated at 76hrs AED, or larvae irradiated with 25 Gy as positive control for cell death (Irradiation). Scale bar = 100μm.

**Figure 1 - supplement 4.**
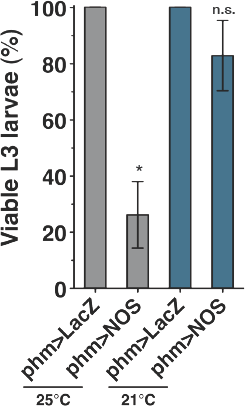
*NOS* overexpression in the PG at 21°C increases larval survival into the 3rd instar. Percent viable L3 *phm>NOS* and control (*phm>LacZ*) larvae raised at 25°C and 21°C. Most *phm>NOS* larvae raised at 25°C die before the third instar (L3). Rearing *phm>NOS* larvae at 21°C increased the number of animals that progress to the third instar. Statistical analysis; mean +/- SEM of three replicates. ^*^ P<0.05 calculated by two-tailed Student’s t-test.

**Figure 2 - supplement 1.**
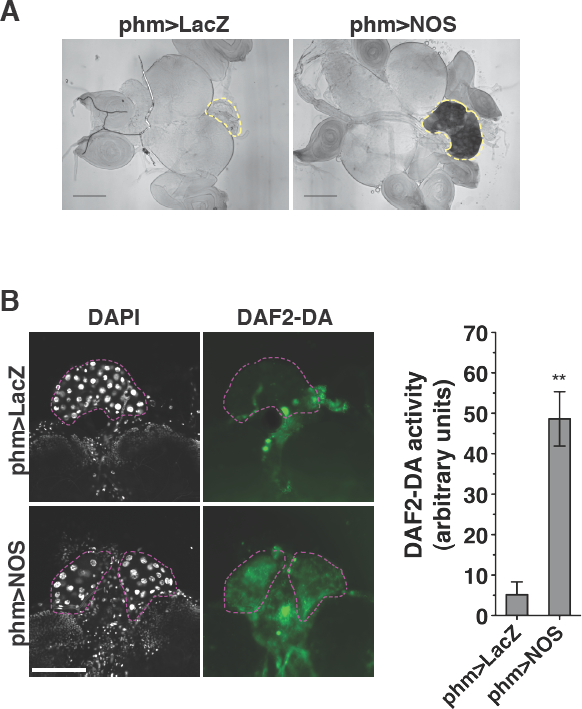
DAF2-DA assay for detection of NO production. (A) NOS enzymatic activity visualized by NADPH-diaphorase staining in targeted overexpression of *NOS* to the PG (outlines) (*phm>NOS*) and control (*phm>LacZ*). Larvae were raised at 21°C and brain complexes were dissected from wandering larvae. Scale bars = 200μm. (B) *NOS* overexpression in the PG (*phm>NOS*) increases NO production in the PG cells (outlined). Measurement of nitric oxide (NO) production by the fluorescent reporter DAF2-DA. Larvae were raised at 21°C and brain complexes with the PG were isolated and stained at 117hrs AED. Scale bar = 100μm. Statistical analysis: mean +/- SEM. ^**^ P<0.01 calculated by two-tailed Student’s t-test.

**Figure 2 - supplement 2.**
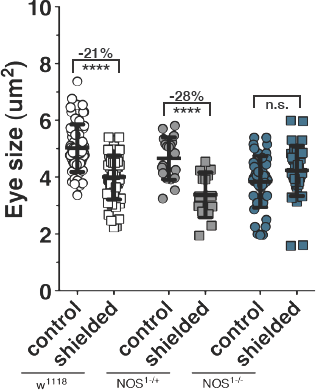
*NOS* is required for coordinating imaginal tissue growth during the regeneration checkpoint. Coordination of growth during shielding irradiation is lost in larvae mutant for *NOS.* Measurement of undamaged eye imaginal disc size following shielded irradiation (25 Gy) compared to unirradiated control (0 Gy) in wildtype (W^1118^) and animals heterozygous or homozygous for *NOS* mutant (*NOS*^1^). Posterior tissues were exposed to 25 Gy ionizing irradiation at 80hrs AED and anterior tissues, including the eye discs, were shielded using lead tape. Statistical analysis: mean +/- SD. ^****^P<0.001 calculated by twotailed Student’s t-test.

**Figure 3 - supplement 1.**
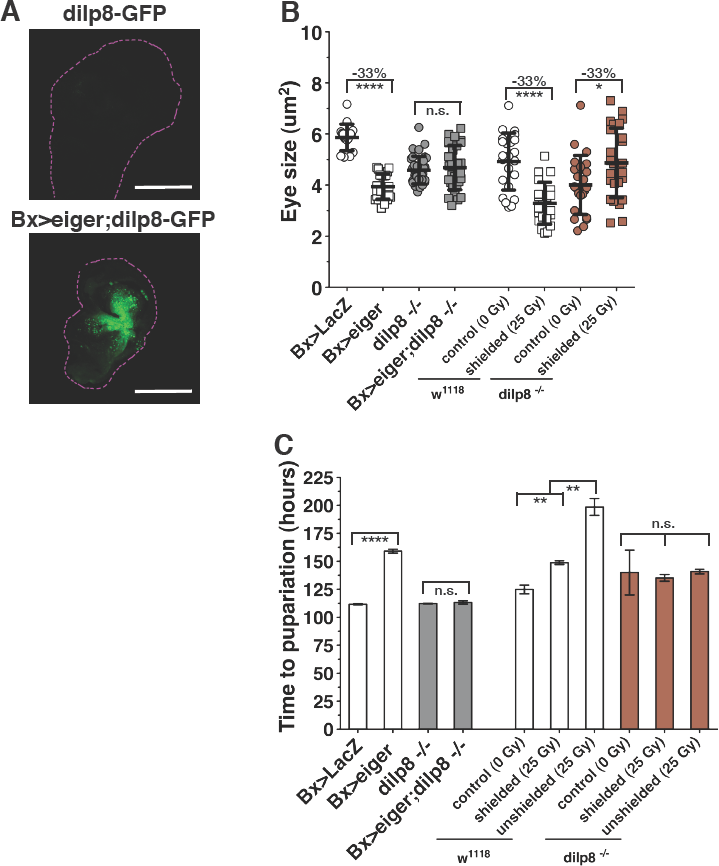
Imaginal disc growth inhibition during eiger-induced damage or shielded irradiation are both dependent on Dilp8. (A) *Dilp8* expression is increased in eiger misexpressing wing imaginal discs. *Dilp8* expression is visualized in control (*dilp8-GFP*) and eiger misexpressing (*Bx>eiger;dilp8-GFP*) wing discs (outlined) using the *dilp8-GFP* enhancer trap (Garelli et al. 2012). Scale bars = 100μm. (B) Eye imaginal disc size measured at 104hrs AED following targeted wing expression of eiger (*Bx>eiger*), or shielded irradiation (shielded) damage, in larvae homozygous for *dilp8*^-/-^ or in WT control larvae. (C) Developmental delay resulting from misexpression of eiger or shielded irradiation is dependent on Dilp8. Measurement of pupariation timing for larvae with targeted wing expression of eiger (*Bx>eiger*) or shielded irradiation (shielded) damage, in larvae homozygous for *dilp8*^-/-^, or in WT control larvae. Statistical analysis: B, mean +/- SD. C, triplicates +/- SEM. ^*^ p<0.05, ^**^ p<0.01, ^****^p<0.001 calculated by two-tailed Student’s t-test, except for shielding experiments in C, calculated by one-way ANOVA with Tukey’s post-test.

**Figure 3 - supplement 2.**
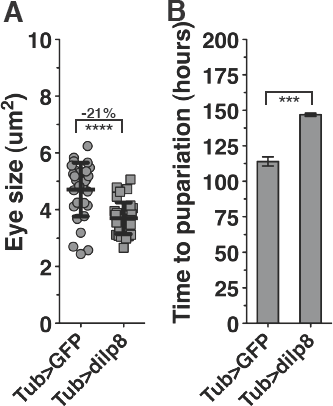
Systemic misexpression of *dilp8* is sufficient to produce developmental delay and imaginal disc growth inhibition. (A) Eye imaginal disc size measured at 104hrs AED following systemic misexpression of *dilp8* (*Tub>dilp8*)or in control larvae (*Tub>GFP*), (B) Measurement of pupariation timing for larvae with systemic (*Tub>dilp8*) misexpression of *dilp8.* Statistical analysis: A, mean +/- SD. B, triplicates +/- SEM. ^***^P<0.005, ^****^P<0.001 calculated by two-tailed Student’s t-test.

**Figure 6 - supplement 1.**
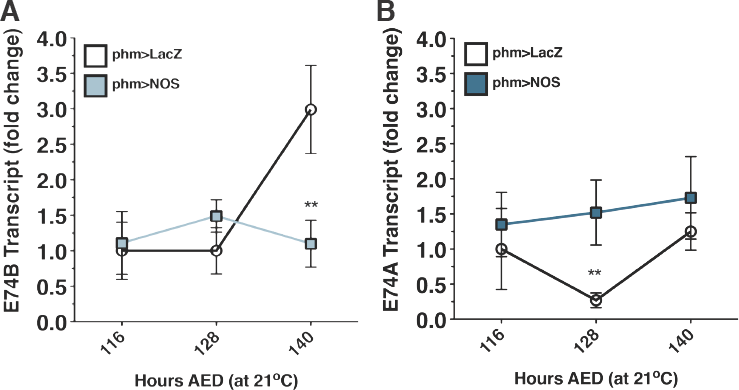
*NOS* overexpression in the PG reduces ecdysone signaling. NOS activity in the PG reduces ecdysone signaling, (A) Expression of *E74B* in control (*phm>LacZ*) larvae and larvae with *NOS* overexpression in the PG (*phm>NOS*), (B) NOS activity in the PG does not precociously induce late-ecdysone signaling, Transcription of ecdysone-induced E74A, a reporter for late ecdysone signaling during the larval-pupal transition, NOS activity in the PG does not increase late-ecdysone signaling, Larvae raised at 21°C. Transcription levels measured by qRT-PCR, normalized to control expression levels at 116hrs AED, Statistical analysis: mean of duplicates +/- SEM. ^**^ P<0.01, calculated by paired one-tailed t-test,

**Figure 7 - supplement 1.**
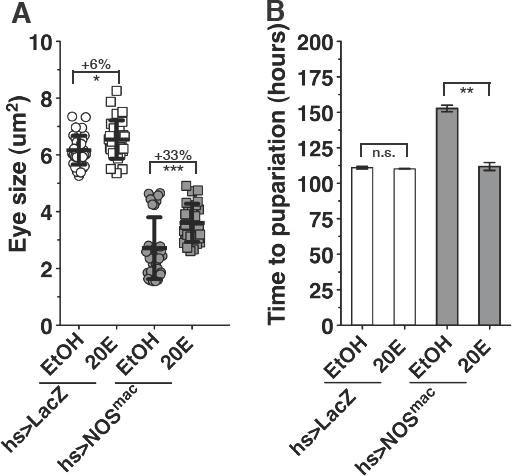
Imaginal disc growth restriction during *NOS*^nac^ misexpression is the result of reduced ecdysone signaling. Ecdysone levels are rate-limiting for Imaginai disc growth and developmental delay during systemic expression of *NOS*^*mac*^ (*hs>NOS*^mac^). (A) 20E feeding rescues growth restriction. (B) Developmental delay induced by *hs>NOS*^mac^ compared to control ethanol only fed larvae (EtOH). Statistical analysis: A, mean +/- SD. B, mean of triplicates +/- SEM. * P<0.05, ^***^P<0.005 calculated by two-tailed Student’s t-test.

**Figure 7 - supplement 2.**
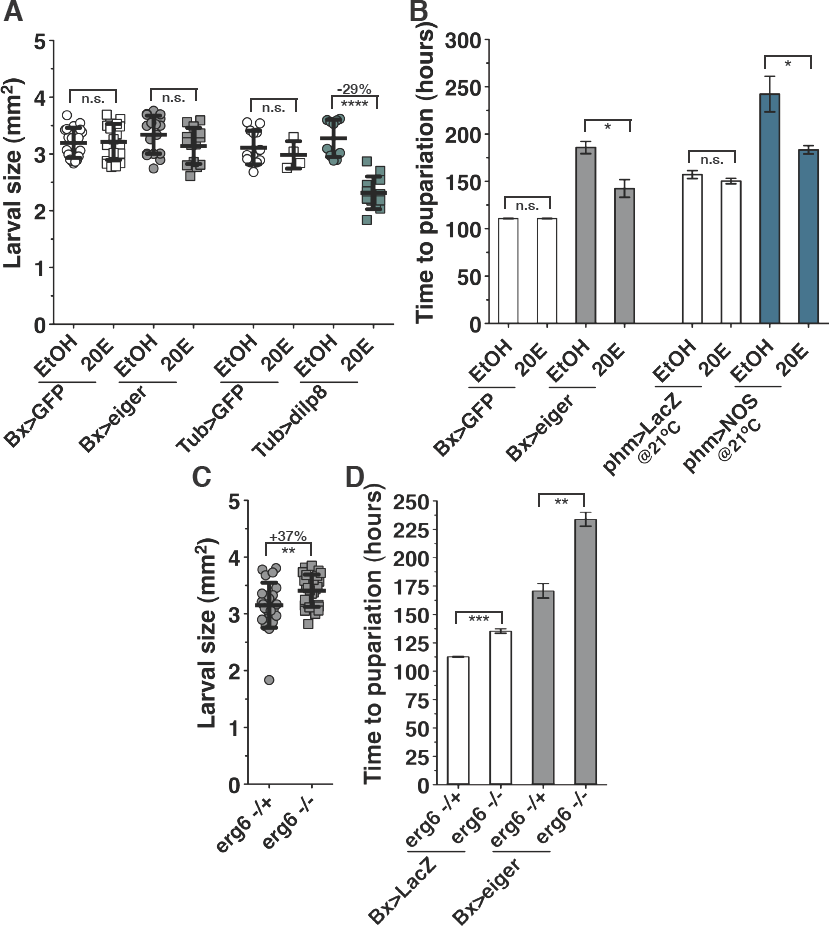
Ecdysone regulates larval growth and developmental time. (A) Measurement of larval growth at 104hrs AED in larvae raised in food with supplemental ecdysone (20E) or control food (EtOH), (B) Measurement of developmental timing to pupariation for larvae raised In food with supplemental ecdysone (20E) or control food (EtOH), (C) *erg*^°^ inhibition of ecdysone increases the growth rate of larval tissues, Measurement of larval growth at 104hrs AED, (D) Restriction of ecdysone synthesis (*erg6*^°^) extends the time to pupation when compared to permissive synthesis conditions (*erg6*^°^+) for both control (*BX>GFP*) and *Bx>eiger.* Statistical analysis: A and D, mean +/- SD. B and C, triplicates +/- SEM. ^*^ P<0.05, ^**^ p<0,01, ^***^P<0.0005, ^****^P<0.001 calculated by two-tailed Student’s t-test.

**Figure 9 - supplement 1.**
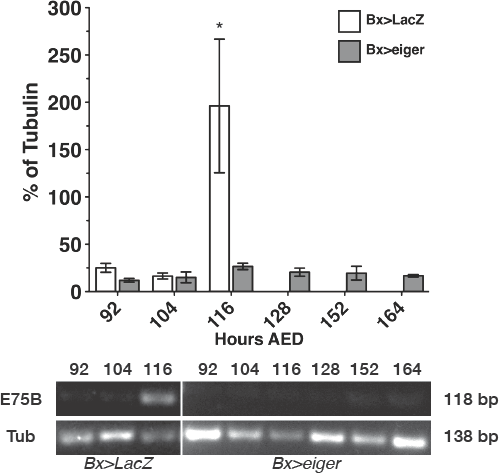
Regeneration checkpoint activation suppresses transcription of *E75B.* Targeted tissue damage (*Bx>eiger*) suppresses transcription of the nuclear hormone receptor (*E75B*) involved in the initiation of pupariation. Transcript levels of *E75B* and tubulin measured by semi-quantitative PCR in *Bx>LacZ* and *Bx>eiger* larvae from 92h AED until the end of the larval growth period. Statistical analysis: mean of two isolation replicates +/- SEM. ^*^ P<0.05, calculated by two-way ANOVA.

**Figure 9 - supplement 2.**
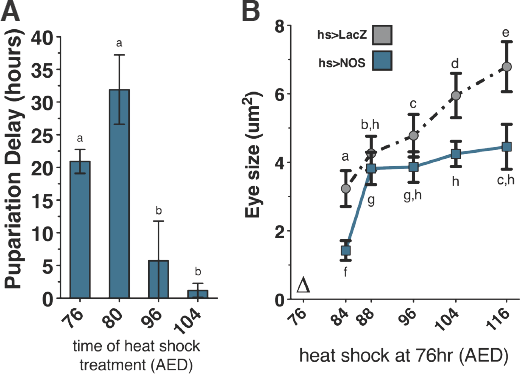
NOS activity during the larval feeding period delays pupation and inhibits growth. (A) Expression of *NOS* during the larval feeding period (76hrs and 80hrs AED) substantially delays larval development, while *NOS* expression during near the cessation of feeding (96hrs and 104hrs AED) does not substantially delay development. *NOS* was systemically expressed by one heat shook treatment (*hs>NOS*) at either 76hrs, 80hrs, 96hrs, or 104hrs AED and time to pupation was measured. (B) Expression of *NOS* during the larval feeding period restricts imaginal disc growth throughout the rest of larval development. *NOS* was systemically expressed by one heat shock treatment (Δ) at 76hrs AED and eye imaginal disc size was measured at subsequent timepoints. Statistical analysis: A, mean +/- SD. B, triplicates +/- SEM. Differing letters denote statistical significance calculated by one-way ANOVA with Tukey’s post-test.

